# Computational Study of Ions and Water Permeation and Transportation Mechanisms of the SARS-CoV-2 Pentameric E Protein Channel

**DOI:** 10.1101/2020.05.17.099143

**Authors:** Yipeng Cao, Rui Yang, Wei Wang, Imshik Lee, Ruiping Zhang, Wenwen Zhang, Jiana Sun, Bo Xu, Xiangfei Meng

**Affiliations:** Tianjin Medical University Cancer Institute and Hospital, National Clinical Research Center for Cancer, Tianjin, 300060 P.R. China; National Supercomputer Center in Tianjin, 300457 P.R. China; Department of Infection and Immunity, Tianjin Union Medical Center, Nankai University Affiliated Hospital. 300031, P.R. China; College of Physics, Nankai University, Tianjin, 300071 P.R. China; Center for Intelligent Oncology, Chongqing University School of Medicine and Chongqing University Cancer Hospital, Chongqing 400030, P.R.China

## Abstract

Coronavirus disease 2019 (COVID-19) is caused by a novel coronavirus (SARS-CoV-2) and represents the causative agent of a potentially fatal disease that is of public health emergency of international concern. Coronaviruses, including SARS-CoV-2, encode an envelope (E) protein, which is a small, hydrophobic membrane protein; the E protein of SARS-CoV-2 has high homology with that of severe acute respiratory syndrome coronavirus. (SARS-CoV) In this study, we provide insights into the function of the SARS-CoV-2 E protein channel and the ion and water permeation mechanisms on the basis of combined in silico methods. Our results suggest that the pentameric E protein promotes the penetration of monovalent ions through the channel. Analysis of the potential mean force (PMF), pore radius and diffusion coefficient reveals that Leu10 and Phe19 are the hydrophobic gates of the channel. In addition, the pore demonstrated a clear wetting/dewetting transition with monovalent cation selectivity under transmembrane voltage, which indicates that it is a hydrophobic voltage-dependent channel. Overall, these results provide structural-basis insights and molecular-dynamic information that are needed to understand the regulatory mechanisms of ion permeability in the pentameric SARS-CoV-2 E protein channel.

## Introduction

COVID-19 is a severe and highly contagious respiratory illness that was first reported in China in early December 2019. Subsequently, the virus has spread worldwide. As of May 6, 2020, millions of cases have been confirmed, and hundreds of thousands have died. The World Health Organization (WHO) announced a global pandemic for COVID-19 in March 2020. In addition to the hazards of the disease itself, it has also led to a severe turbulence in international financial markets and may cause serious consequences such as a financial crisis. COVID-19 is a disease caused by a new coronavirus named SARS-CoV-2. It is speculated that it originated from bats and was transmitted to humans through an intermediate host (some kind of wildlife). Its symptoms include fever, general malaise, dry cough, shortness of breath, and respiratory distress. Comparing similar diseases, including severe acute respiratory syndrome (SARS) and Middle East respiratory syndrome (MERS), which are also caused by coronaviruses, the mortality rates for SARS and MERS were 10% and 36%, respectively ^1^. Currently, COVID-19 has a mortality rate of 1 to 10% in different countries but appears to be more contagious than SARS and MERS ^2 3 4 5^.

Similar to other coronaviruses, SARS-CoV-2 is a long positive-sense, long single-stranded (30 kb) RNA virus. The structure of different human coronaviruses (HCoVs) is similar. The viral genome is packed by nucleocapsid (N) proteins, forming a helicoidal nucleocapsid protected by a lipid envelope ^6^. Several viral proteins, including the spike (S), envelope (E), and membrane (M) proteins, are embedded within a lipid envelope ^7 8^. Studies have shown that S, M, and N proteins play important roles in receptor binding and virion budding. For example, the M protein participates in virus germination and interacts with the N and S proteins. The S protein has immune recognition sites and can be used to design vaccines ^9^.

Currently, the importance of the E protein has not been fully revealed. Evidence suggests that the E protein maintains its morphology after virus assembly by interacting with the M protein ^10 11 12^. When an E protein gene mutation occurs, it promotes apoptosis. Recent studies have shown that coronaviruses have a viroporin that can self-assemble into a pentameric structure and have ion selectivity. When there is a transmembrane voltage, the ion channel characteristics of viroporins are more significant. In addition, the Asn25Ala and Val18Phe mutations could destroy ion channel activity ^13^. This indicates that the E protein may play an important role in regulating the ion equilibrium inside and outside the viral envelope.

The ion channel activity of the E protein can lead to increased levels of the inflammatory cytokines IL-1β, TNF and IL-6 in the lungs, leading to the occurrence of an “inflammatory storm” ^14^. This plays a key role in the progression of the disease and may cause the patient’s condition to suddenly deteriorate and lead to death. In addition, its evolutionary conservatism may be an important cause of viral cross-host infection ^15 16 17^.

Although previous studies have suggested that the E protein of coronaviruses such as SARS and MERS oligomerizes and has ion permeability, the specific mechanism of ion permeability and channel properties remain to be explored due to the lack of a crystal structure for the E protein. In 2014, Li et al ^17^ identified the SARS-CoV E protein monomer structure. In 2018, Surya et al. extracted the pentamer structure of the SARS E protein by nuclear magnetic resonance (NMR) ^18^, which provided strong support for the study of the SARS-CoV-2 E protein pentamer ion permeability mechanism.

In this study, we obtained the amino acid sequence of the SARS-CoV-2 E protein from the National Center for Biotechnology Information (NCBI) database ^19^. The E protein pentamer model of SARS-CoV-2 was built by using the homology modeling method, and the reasonableness of the model was evaluated. Subsequently, µs -level molecular dynamics (MD) simulations were performed to evaluate the pentamer’s stability in the membrane environment. We tried to use potential mean force (PMF) to reveal the permeability of different physiological ions and water molecules in the pores of the E protein pentamer. The characteristics of the pentameric channel were analyzed by combining the channel diffusion coefficient and geometric properties. In addition, computational electrophysiology was applied for different transmembrane voltages of the system to reveal the effect of the voltage on the ion permeability of the pentameric E protein.

Overall, exploring the mechanisms of the pentameric SARS-CoV-2 E protein not only provides valuable insights into the conduction of the channel but also has important implications for our understanding of the difference between SARS-CoV-2 and other coronaviruses.

## Results

### Sequence alignment and homology modeling

Figure 1A shows the sequence alignment between the E proteins of two human coronaviruses (SARS-CoV and SARS-CoV-2 sequence) made by Clustal X software. The blue dotted rectangle in Figure 1B is the area of SARS-CoV-2 that corresponds to the crystal structure of the SARS-CoV E protein. The E protein amino acid sequences of these two coronaviruses are highly homologous, and the overall and transmembrane (TM) region similarity reached 91.8% and 100%, respectively.

**Figure 1.**
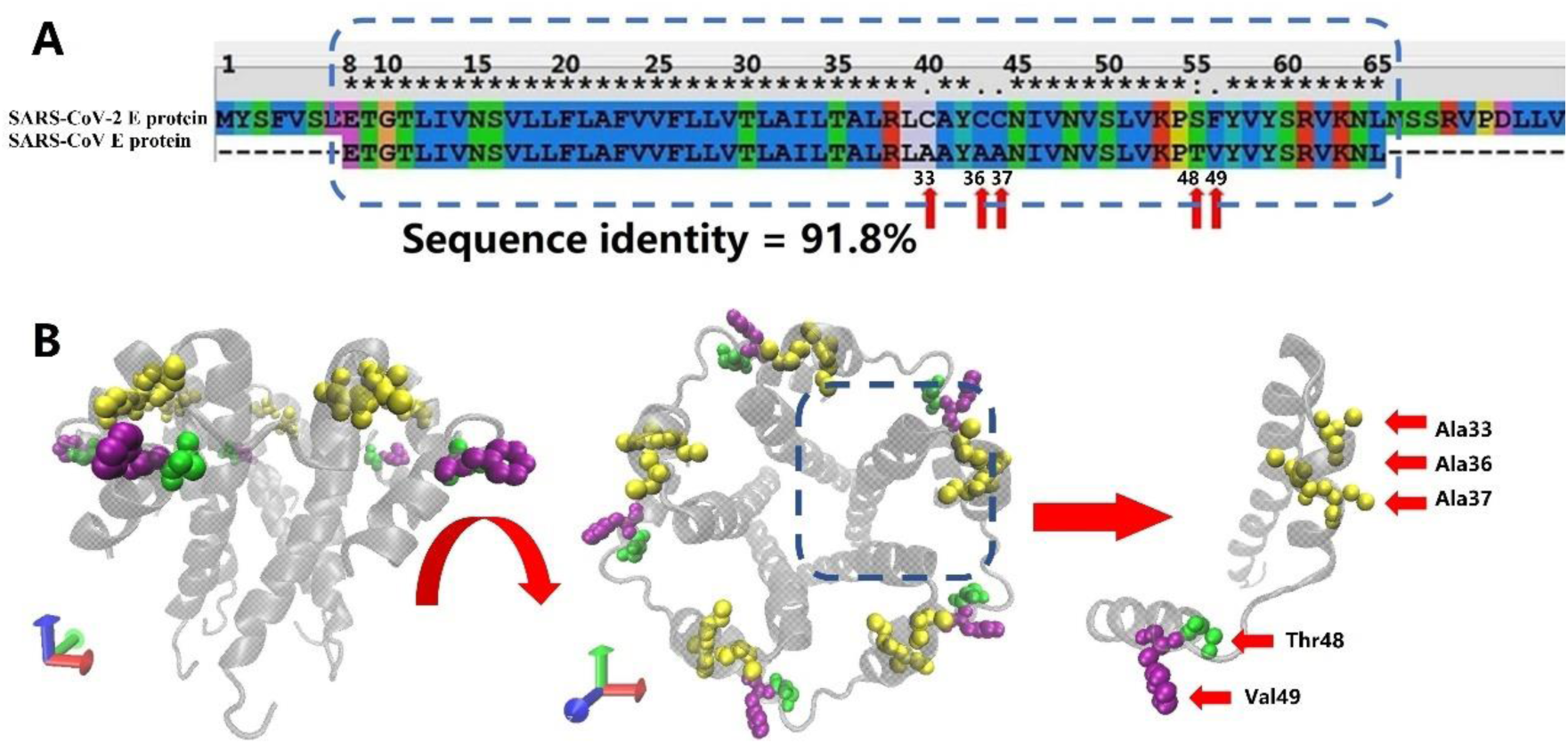
A, The sequence alignment for the E protein of SARS-CoV-2 and SARS-CoV. The SARS-CoV E protein sequence is from the protein data bank (PDB) database (PDB ID: 5X29). The SARS-CoV-2 E protein sequence was obtained from NCBI. The sequences in the rectangular box are used for homology modeling. The red arrows mark the different amino acids between the SARS-CoV-2 and SARS-CoV E proteins. B, Two views of the E protein model of SARS-CoV-2. The different amino acids between SARS-CoV and SARS-CoV-2 are marked with red arrows.

### SARS-CoV-2 E protein pentamer homology model evaluation

The Rampage online program ^20^ was used to evaluate the accuracy of the SARS-CoV-2 pentameric E protein model. More than 98.9% of the amino acids are within the acceptable range, suggesting that the SARS-CoV-2 E protein is similar to the SARS E protein NMR model. Subsequently, a 1000 ns (1 µs) MD simulation was performed to evaluate the stability of pentameric E protein embedded in the membrane environment. The root-mean-square deviation (RMSD) of the model is shown in Figure 2, and the red and blue curves represent the whole protein and TM region, respectively. In the first 200 ns, the RMSD continued to rise, indicating that the model needed longer optimization (compared with other ion channels) to reach the pentameric structure equilibrium. During the last 800 ns, the curves plateaued. The RMSD of the whole protein and the TM region converged at ∼0.4 and ∼0.3 nm, respectively. There is a 0.1 nm difference between the whole protein and TM region. These findings are consistent with other ion channel data that show that the TM region has a higher stability than the other parts of the membrane protein ^21^.

**Figure 2.**
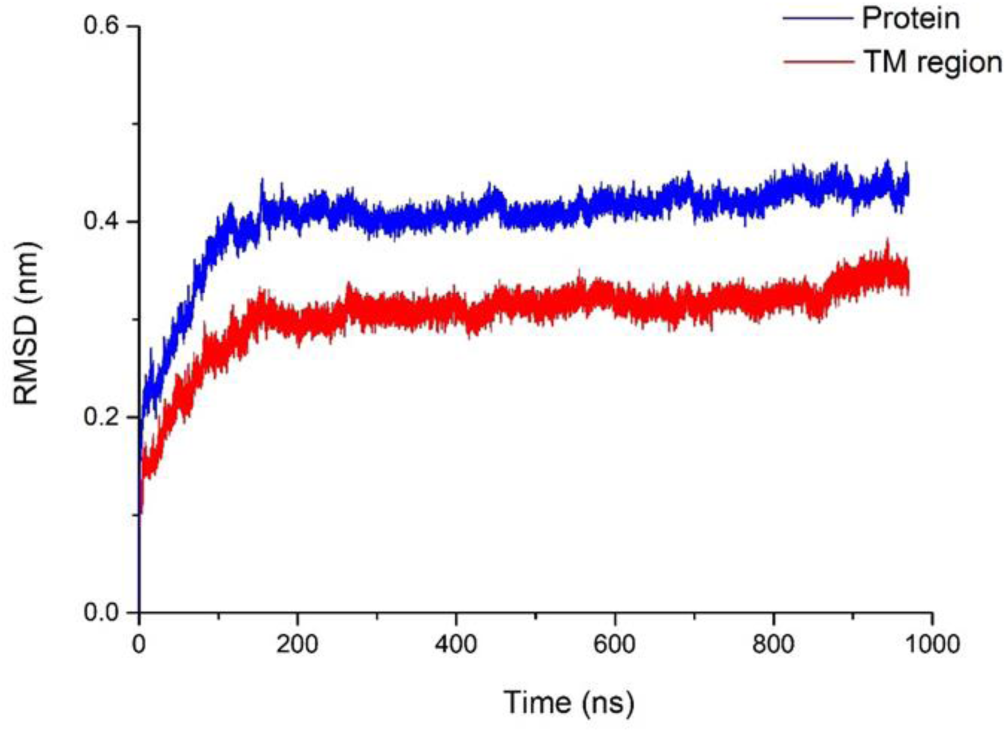
Structural stability of the SARS-CoV-2 E protein. The blue and red curves represent the root-mean-square deviation (RMSD) of the whole pentameric E protein and TM region, respectively.

### Water and ion permeability

The permeability of the SARS-CoV-2 pentameric E protein channel is very important for understanding the replication ability of viruses in cells and how they are secreted into the extracellular medium ^22^. It is possible to examine a profile of the free energy of a single ion or water as a function of its position along the pore axis by calculation of PMF. In this simulation, the ions or water molecules are restrained to a continuous position along the z-axis and move freely in the pore’s xy plane. Moreover, other parts of the system (proteins, ions, water molecules, and lipids) can move freely and reach equilibrium.

The PMF of important physiological ions (Mg2^+^; Ca^2+^; Cl-; K^+^ and Na^+^) and the water molecules as a function of their position along the pore (z)-axis were calculated separately. Figure 3A shows the PMF of ions and water molecules permeating through the SARS-CoV-2 E protein pentamer pore. Z_M_ is defined as the axial direction along the pore, which is from −3 ∼ 3 nm (the length of umbrella sampling is 6 nm in total). From the PMF curve, we found that the energy barriers of monovalent and divalent ions have a significant difference. The maximum energy barriers of the two divalent ions Ca^2+^ and Mg^2+^ are 60 kJ/mol and 70 kJ/mol, respectively. In contrast, the PMFs of the monovalent ions Na^+^ and K^+^ are ∼ 12 kJ/mol and ∼20 kJ/mol, respectively, while the Cl-energy barrier is ∼ 30 kJ/mol, which is significantly smaller than that of Ca^2+^ and Mg^2+^. From an energy perspective, the SARS-CoV-2 pentameric E protein is almost impermeable to divalent ions. This confirms the previous hypothesis that the SARS-CoV and MERS-CoV E protein channel is a monovalent cation channel ^23 24^. The maximum energy barrier in descending order is as follows: Na^+^ <K^+^ <Cl-<Mg^2+^ <Ca^2+^. Additionally, the barrier of water molecules is only ∼ 5 k/mol, which is similar to the energy threshold of free diffusion in liquid. Interestingly, there is no obvious peak in the PMF curve for Na^+^ ions and water molecules. This result indicates that the E protein pentamer pores are not permeable to divalent ions. The difference in the energy barrier between the monovalent cations is large, suggesting that the channel has selective permeability for specific ions.

**Figure 3.**
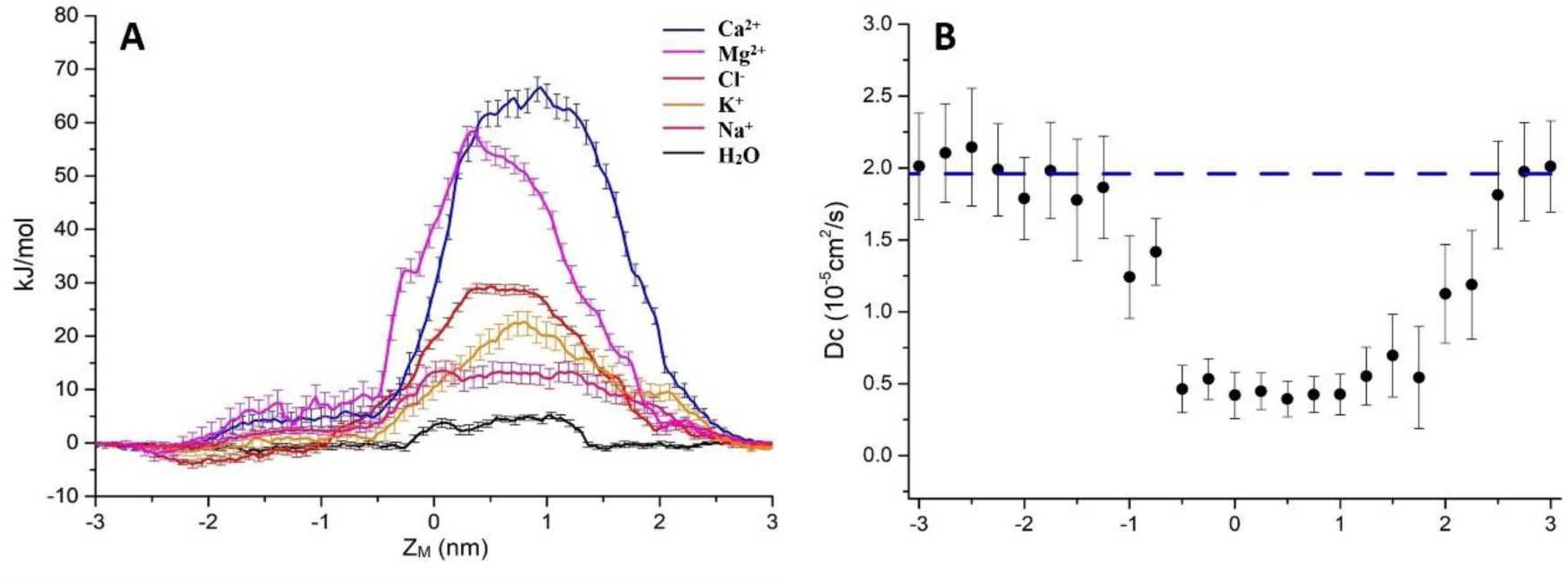
A, PMF profiles of single ions (Ca^2+^, Mg^2+^, Cl^-^, K^+^, Na^+^, H_2_O represented by blue, magenta, red, yellow, pink and black, respectively) as a function of position along the pentameric SARS-CoV-2 E protein pore axis. B, The diffusion coefficient profile. The dashed line is the experimental value of a single K^+^ diffusion coefficient in water, D_C_=10^−5^ cm^2^/s. Error bars were estimated by bootstrapping and are the same color as the corresponding line.

To explore the permeability of the channel, we calculated the diffusion coefficient (D_C_) using a single K^+^ ion as a probe. By fitting the K^+^ mean square displacement (MSD) with the Einstein equation (MSD = 2D(c)t), we obtained the diffusion coefficient. As shown in Figure 3B, the average diffusion coefficient of K^+^ outside the pore is ∼ 2.0×10^−5^±0.4 cm^2^/s, which is consistent with the experimental data of K^+^ in the liquid environment ^25^. However, the D_C_ decreases by approximately 70% when the K^+^ probe is in the channel pore. The D_C_ curve of the internal channel pore has a flat bottom and is similar to the PMF. The mid-value of D_C_ = 0.35×10 ^-5^±0.2 cm^2^/s indicates that there is a large obstacle for probe diffusion in the inner pore, showing a similar value to other narrow hydrophobic pores ^26 27^.

Next, we used the solubility-diffusion equation (Eq 1) ^28 29^ to evaluate the permeability coefficient (P) between two different ions.

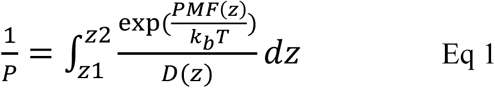

In Eq 1, PMF(z) and D(z) are functions of PMF and DC along the z-axis, z1 denotes a position on one side of the pore and z2 denotes a position on the other side. Kb is the Boltzmann constant, and T is the simulation temperature. The part of the PMF in the TM region is approximate to the graph of the quadratic function, so the PMF(Z) uses quadratic function fitting. The linear fitting of the corresponding D_C_ gives the function D(z). The ratios of the P for Na^+^, K^+^ and Cl-in the maximum energy barrier region were simply evaluated. P_Na+_/P_K+_ = 22.42, P_K+_/P_Cl-_ = 5.82, and P_H2O_/P_Na+_ = 402.57.

### Computational electrophysiology

Previous electrochemical experiments have shown that the transmembrane voltage has an important effect on the pore permeability for the E protein in both SARS-CoV and MERS-CoV ^23 24^. In addition, viroporins and similarly nanopores are sensitive to transmembrane voltage, causing electrowetting of the pores and changing the tension on the hydrophobic surface inside the pores, which is enough to functionally open the channels ^30 31 32 33^. We used the computational electrophysiology first proposed by Kutzner et al. This approach expands the membrane protein structure into a “sandwich” and simulates the transmembrane voltage by adjusting the number of ions at different layers. It can directly simulate and analyze the ion penetration in the presence of transmembrane voltage at an atomic resolution. The sandwich structure of the SARS-CoV-2 E protein pentamer is presented in Figure 4A. A total of 4 models were established to simulate the charge differences between the two sides of the channel (ΔQ = 12e, 8e, 4e, and 0e). Using the method described above, the transmembrane voltage was calculated as ΔU = ∼ 0.45 V, 0.3 V, 0.15 V, and 0 V, respectively (Fig 4B).

**Figure 4.**
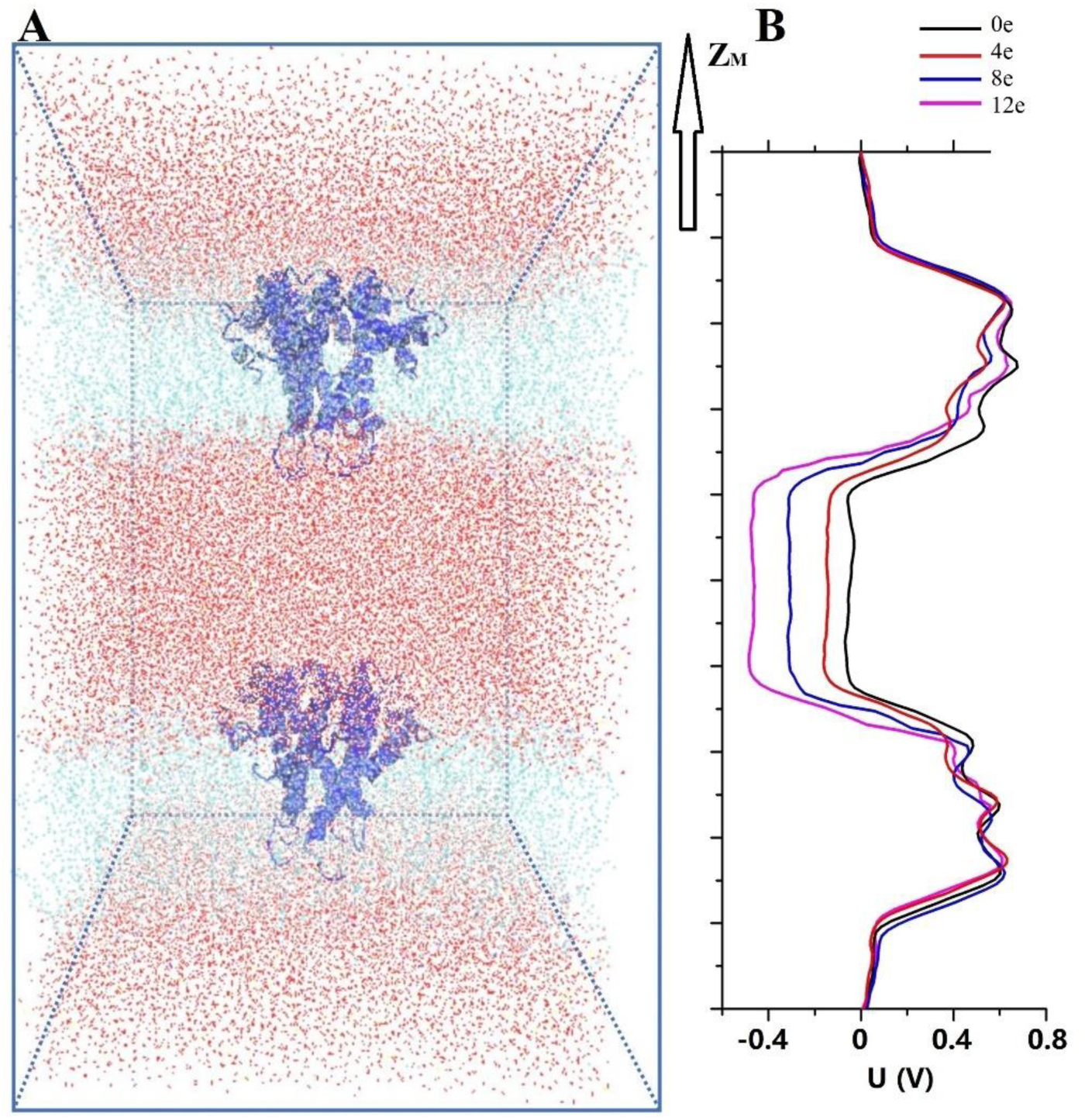
A, A sandwich simulation system of SARS-CoV-2 at 0.15 M NaCl concentration, and this includes the SARS-CoV-2 pentameric E proteins, cell membrane, ions and water molecules. B, Average time of the electrostatic potential ΔU along the z-axis, arising from imbalances ΔQ between the elementary charges of 0 and 12(color curve).

### Effect of transmembrane voltage

Trick et al. indicated that the behavior of water molecules in pores can be used as a reasonable method to study the ion permeability of channels ^27^. Many works have shown that water influx into hydrophobic pores may help ions overcome the energy barrier on hydrophobic surfaces ^34 35^. Overall, we could speculate the ion penetrability of the E protein pentamer by the statistical number of water molecular counts in the inner pore.

To accurately assess the behavior of water molecules in the SARS-CoV-2 E protein pentamer pores, we performed six 20 ns simulations for each transmembrane voltage, and the total simulation time reached ∼0.5 µs. Subsequently, the water molecules at different transmembrane voltages were counted (the statistical data of the changes in the molecule counts are shown in the Supplementary Word Film). We extracted representative water molecule diagrams for each voltage, as shown in Figure 5A. When ΔQ = 0e, the average time for the channel to keep wetting is ∼ 3 ns. ΔQ = 4e is slightly longer (∼ 5 ns). The absolute number of water molecules in the pore is small, with the average ranging from 0 to 5 (Fig 5B). Although the pores are functionally open for water permeation, due to the small stream of water, they may not be able to maintain sufficient electrical wetting, resulting in ions that cannot permeate unimpeded. We speculate that at the maximum physiological voltage (∼0.2 V), the SARS-CoV-2 pentameric E protein pore can only permeate a few water molecules.

**Figure 5.**
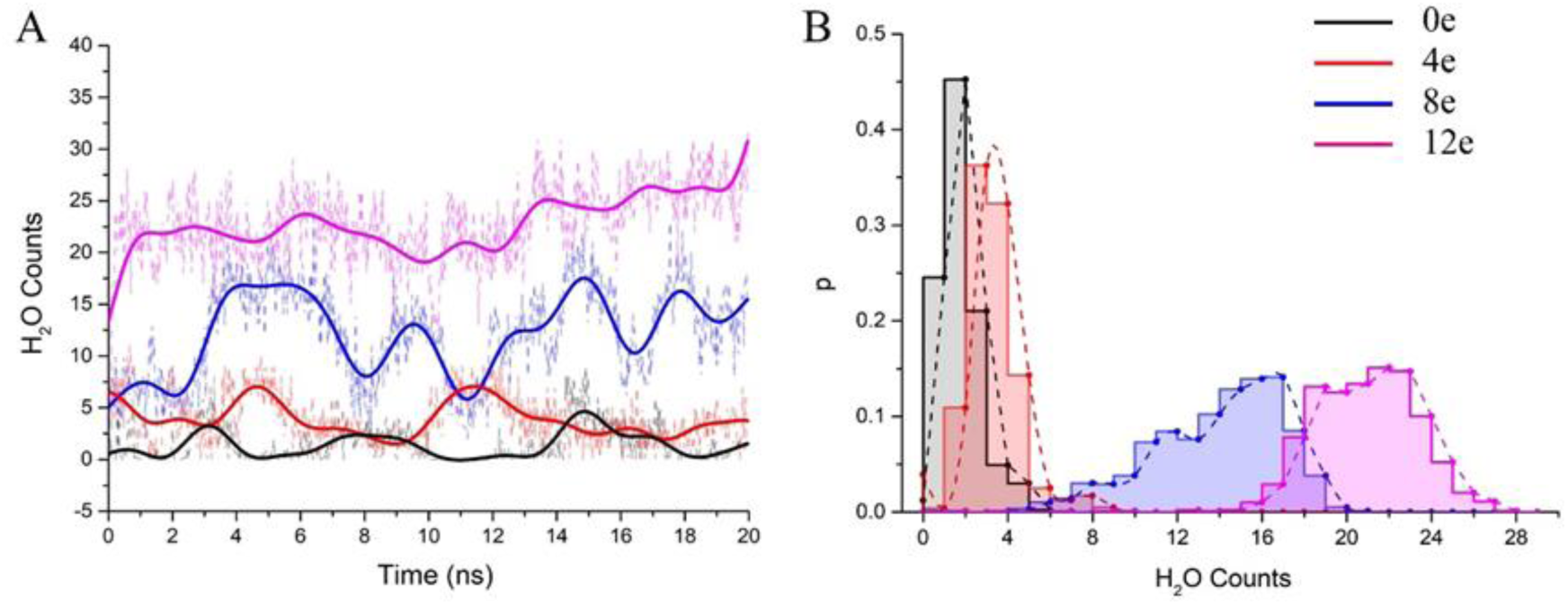
A, The change in water molecule counts in the inner pore of the channel under different transmembrane voltages. The scatter represents the absolute counts of water molecules, and the color curve represents after smoothing (n = 25) using the Fast Fourier Fransformation (FFT) method. B, Normalized probability histogram of the water counts in the TM region for 20 ns simulations. The different colors represent different models: 0e (blue), 4e (red), 8e (blue) and 12e (magenta).

The number of water molecules increased significantly when ΔQ = 8e or 12e (ΔU = ∼ 0.3 V, 0.45 V), and the water distributions were 4-20 (8e) and 15-27 (12e) (The process of water permeation see supplementary Movie 1). The pores remained in a wetting state. The experiment shows that the maximum ΔU = 0.45 V is not enough to cause electrical breakdown of the lipid bilayer ^36^, indicating that the SARS-CoV-2 E protein channel is activated by transmembrane voltage. Additionally, we observed that a few water molecules were able to be cotransported with Na^+^ and K^+^ during the simulations (See Supplementary Movie 2 and 3). Interestingly, as the voltage increased, the permeability of water molecules also increased synchronously, showing a periodic change regardless of the magnitude of the transmembrane voltage (even if ΔU = 0). The calculation suggests that the SARS-CoV-2 E protein pentamer channel has obvious wetting/dewetting transitions and that it has the typical characteristics of voltage-dependent hydrophobic channels.

### Pore conformation analysis

By calculating the channel radius at different voltages (ΔU = ∼ 0.45 V, 0.3 V, 0.15 V, 0 V and 0 V (closed state)) (Fig 6B), we found that in the pentameric E protein channel in the closed state (ΔU = 0 V), there are two positions with the minimum radius in the TM region, r = 0.2 nm (Z_M_ = −0.8 nm) and 0.15 nm (Z_M_ = −0.5 nm), corresponding to Leu10 and Phe19 (blue and red arrows, respectively). Phe19 and Leu10 are located at the top and bottom of the pore, respectively. The two positions show a candidate gated area of the channel. Figure 6A shows that these two locations have obvious “bottlenecks”, which are not large enough for water permeation. This was also confirmed by the calculation of the channel radius on different transmembrane voltages. The radius of the pore obviously increased with transmembrane voltage at the two locations. In the open state (ΔU=0 V), the pore radius increased to 2 nm and 2.2 nm. When the transmembrane voltage increased to ΔU = ∼ 0.15 V and 0.3 V, the radius of the pore at the Leu10 position was ∼ 0.23 nm and 0.28 nm, and the radius of the pore at Phe19 was ∼0.25 nm and 0.33 nm, respectively. The voltage had a greater influence on Phe19 than on Leu10. However, for ΔU = 0.45 V, the radius of the pore was still ∼ 0.33 nm at Phe19 but had expanded to 0.4 nm at Leu10, indicating that the large transmembrane voltage may make Leu10 excessively available, which may cause the unidirectional permeation to be destroyed. The narrowest locations of the inner pore are consistent with the energy barrier. Apparently, the radius of the inner pore showed that Leu10 and Phe19 of the SARS-CoV-2 E protein are candidate hydrophobic gates.

**Figure 6.**
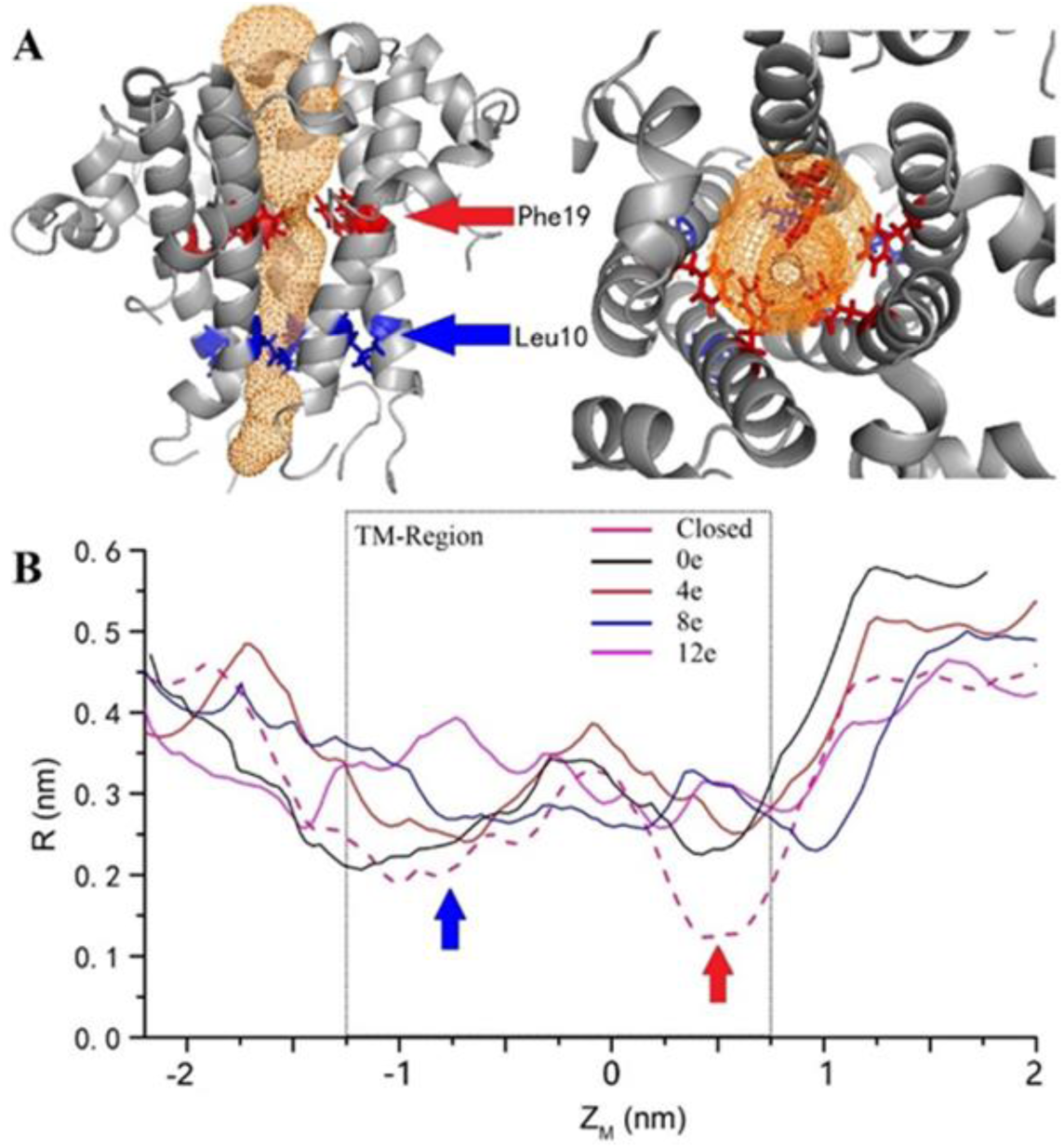
A, The channel structure in the closed state. The red and blue arrows indicate the two residues with the largest radius changes at different voltages. B, The variation in the inner pore radius. The z axis shows the distance from the center of mass. The black, red, blue, and magenta solid curves represent ΔQ=0e, 4e, 8e, and 12e, respectively, and the pink dashed curve represents the closed state. The calculated channel as measured by CAVER 3.01 ^37^.

## Discussion

COVID-19 is an unprecedented threat to humanity. To understand the infection, the replication mechanism of the virus in the human body is particularly important. Currently, research on the pathogenic mechanism and infection route of COVID-19 mainly focuses on the S protein ^38 39^, but research on the E protein is lacking. Previous reports note that the E protein pentamers from different coronavirus subtypes (SARS-CoV, MERS-CoV) may be viroporins, which play an important role in the virus life cycle ^40^. Deleting the encoding gene of the E protein will greatly reduce the replication and pathogenicity of coronaviruses. Moreover, the pentameric E protein channel activity is necessary for inflammasome activation and is the determinant of coronavirus virulence ^41^. This study explored the permeability mechanism of the SARS-CoV-2 pentameric E protein. We used the NMR conformation of the SARS-CoV E protein pentamer to build the SARS-CoV-2 E protein pentamer model by homology modeling. A combination of MD simulations, PMF, diffusion coefficient calculations, computational electrophysiology analyses, and channel pore geometry analyses was used. The results were mutually corroborated and characterized the TM dynamics of the SARS-CoV-2 pentameric E protein channel. It was found that the pentamer is a monovalent cation-selective voltage-dependent hydrophobic channel, and the two amino acids Leu10 and Phe19 are candidate hydrophobic gates.

The SARS-CoV-2 E protein pentamer has been considered an unknown nature ion channel. There is significant similarity in the TM region sequences (reaching 91.8%) and high gene homology between the SARS-CoV and SARS-CoV-2 E proteins; five different residues are all located at the region of interaction between the protein and membrane, while the sequences of the TM region are completely identical. The highly conserved TM region also suggests similar E protein functions in different coronavirus subtypes. As the RMSD of the E protein pentamer during the 1 µs MD simulation, the structure has a certain flexibility, showing that its geometric deformation can make the channel permeable to ions and water molecules. Figure 2 shows that the TM region of the SARS-CoV-2 E protein remains relatively stable during the simulation, indicating the stability of the pentamer in the membrane environment, which is consistent with previous studies of SARS-CoV and MERS-CoV.

The free energy calculation of the ions permeating through the SARS-CoV-2 pentameric E protein channel strongly suggests that the pore has selection permeability for monovalent ions. The maximum energy barriers of Na^+^, K^+^, and Cl^-^ are 15 kJ/mol, 20 kJ/mol, and 30 kJ/mol, respectively. The comparison of the P values for Na^+^, K^+^ and Cl^-^ shows that the P of Na^+^ is 22 times greater than that of K^+^, while the P of K^+^ is 6 times greater than that of Cl^-^. Although the computational permeability coefficient is two times higher or lower than that observed experimentally for SARS-CoV, it still shows a very large gap in the permeation between ions and water molecules. Additionally, there is no obvious peak in the PMF when Na^+^ and water molecules permeate through the channel, further supporting that Na^+^ and water molecules have less penetration resistance at the inner pore. This further confirmed the selectively for Na^+^. The ion selectivity of SARS-CoV-2 is different from that of SARS-CoV ^23^. Because the maximum energy barrier of Mg^2+^ and Ca^2+^ is 2 ∼ 3 times higher than that of the monovalent ions, divalent ions cannot overcome such a very large energy barrier under the simulation environment. The conformational analysis indicates that it is mainly caused by the geometric size of the pore (Fig 6).

The importance of transmembrane voltage on viroporin has been found in many viruses, including HIV1 Vpu oligomers and the hepatitis C virus (HCV) p7 protein, etc. ^42 43 44^. Although it has been observed in many experiments, this mechanism has not been revealed. Because water molecules play an important role in hydrophobic ion channels, their behavior at the inner pores under different voltages (ΔU =∼0.45 V, 0.3 V, 0.15 V, and 0 V) is the focus of this simulation. Computational electrophysiology enabled the determination of a conducive conformation for ions. The channel wetting/dewetting transitions appeared under different transmembrane voltages, and the radius of the pore increased. Meanwhile, the capacity to contain water molecules also increased. Further analysis of the trajectories revealed that the monovalent cations (Na^+^ and K^+^) will permeate through the pore with water molecules when the transmembrane voltage is > 0.3 V.

Computational electrophysiology also explained that the transmembrane voltage led to an easier wetting transition for the channel. The range for keeping the channel open should be between 0.15 V and2 0.45 V. Na^+^ and K^+^ could be cotransported during water permeation, which is very similar to that in many hydrophobic channels ^45 46 47 48^. Intriguingly, we did not observe Cl^-^ conduction, which may be because chloride ions could not overcome the energy barrier under the maximum transmembrane voltage. In addition, the change in the pore radius at different voltages highlights two important sites, where the geometric radius changes the most, corresponding to Leu10 and Phe19. Phe19 contains a benzene ring group at the hydrophobic surface of the inner pore as a hydrophobic gate. The isopropyl group of Leu10 at the bottom of the pore prevents the reverse penetration (Fig 6A). Previous studies indicated that Asn15Val and Val25Phe mutations will make the channel dysfunctional. This may be due to the additional side chain groups in the pores caused by these mutations (especially the benzene ring in Val25Phe), making 1) the radius of the pores decrease and 2) the hydrophobicity of the pores increase. In short, these mutations may increase the energy barrier, negatively affecting the wetting transition and causing the channel to be functionally closed.

Therefore, the SARS-CoV-2 E protein pentamer is a voltage-dependent hydrophobic channel. We propose that the E protein may play an essential role in the virus infection and replication processes through the following mechanisms: 1) The monovalent selective permeation of the E protein pentamer ion channel may change the intracellular pH, providing a suitable microenvironment for virus replication. 2) Selective permeation can form a transmembrane voltage, creating feedback regulation and maintaining an intracellular microenvironment that is suitable for viral growth. 3) The disintegration of the ion equilibrium of the intracellular area affect the charges coming from the cell through ion channels in the cell membrane, changing the pH and making it easier for the virus to fuse with the cell membrane. This is the infection mechanism of the avian coronavirus as well as some types of influenza viruses ^49 50^. 4) It has a certain signal transduction function.

## Conclusion

Although the pentameric SARS-CoV-2 E protein crystal structure is not yet available, the channel conformation was built by homology modeling from the SARS-CoV E protein pentamer. We characterized and estimated the channel’s stability, energy barrier, voltage dependence and geometric properties by using MD simulations. Our simulation results demonstrate that the SARS-CoV-2 E protein pentamer is a voltage-dependent hydrophobic channel with monovalent cation selectivity. Water molecules and monovalent cations can penetrate through the channel spontaneously under a transmembrane voltage. Differences in the penetrability of E protein channels may be responsible for the infectivity, inflammatory response and incubation period of SARS-CoV-2, which is different from that of SARS-CoV and MERS-CoV. The results further suggested that Leu10 and Phe19 are hydrophobic gates for ion regulation and permeability. Our research reveals the nature of the SARS-CoV-2 E protein pentamer, providing theoretical support for future research on vaccines against E proteins from SARS-CoV-2 and other coronaviruses.

### Experimental methods

Preparation of the SARS-CoV-2 pentameric E protein-membrane simulation system The amino acids sequence of the human coronaviruses E proteins were downloaded from the National Center for Biotechnology Information (NCBI). The Cluster-X software was used for Multiple Sequence Alignment (MSA) for the new human coronavirus and the SARS-CoV E protein. The model was built for the SARS-CoV-2 E protein using the SWISSMODEL web server ^51^. As the SARS E protein (PDB ID: 5X29) ^18^ pentameric structure is the closest E protein to SARS-CoV-2, it was used as a template for building the SARS-CoV-2 E protein. The Rampage web tool was used to evaluate the rationality of the SARS-CoV-2 E protein model. Then, the model was inserted in a DOPC/POPC,1:1 lipid bilayer via pre-equilibrium. The simulation system was set as a 11×11×12 nm^3^ cubic box by CHARMM-GUI ^52^, comprising the whole pentameric E-protein, 330 lipid molecules, ∼30,000 TIP3P water molecules, and concentration is 0.15 M Na^+^ and Cl^-^ ions. Resulting in a simulation system size of ∼120,000 atoms.

#### MD Simulaiton

To obtain the equilibrated pentameric SARS-CoV-2 E protein model, Charmm36 all-atom force field ^53^ was chosen for MD simulation. The MD time step was set at 2 fs. Electrostatic interactions were described using the Particle Mesh Ewald (PME) algorithm ^54^ with a cut-off of 1.2 nm. The LINCS algorithm ^55^ was used to constrain the bond lengths. The pressure was maintained semi-isotropically at 1 bar at both x and y directions using the Perinello-Rahman barostat algorithm ^56^ and the system temperature was maintained at 310 K by the Nose-Hoover thermostat ^57^. Then, a 1000 ns (1 µs) MD simulations were performed for the E protein pentameric.

#### Umbrella sampling

The initial system for umbrella sampling simulations was derived from the equilibrated pentameric SARS-CoV-2 E protein mentioned above. A single ion that maintain physiological activity (Na^+^, K^+^, Ca^2+^, Mg^2+^, Cl^-^) or water molecule was placed at successive positions along the central pore axis by using GROMACS pull code. Energy minimization was performed before simulation for optimizing the water and ions position. The reaction coordinate defined from z +3 to −3 nm with the mass center at z = 0 nm, with a spacing of 0.1 nm between successive windows, resulting in 60 umbrella sampling simulation systems. The probe ion or water molecules were harmonically restrained by a force constant of 2000 kJ mol/nm^2^ same with the pore direction. Each window was performed a 5 ns total umbrella sampling simulation. The initial 2 ns for system equilibration, then a subsequent 3 ns was applied for analysis. The PMFs were computed by the weighted histogram analysis method (WHAM) ^58^, and the profile was generated by Gromacs protocol ‘g_wham’ ^59^. Bootstrap analysis (N = 50) was used to estimate statistical error and the ‘-cycl’ parameter was used to make the value equal.

#### Diffusion-coefficient calculation

The Diffusion-coefficient was calculated using the method described by Shirvanyants et al. ^60^. We restrained the E-protein pentamer helix backbones and made the side chains, membrane, water and ions free to move. Due to the ion selectivity is related to the side chain of inner pore ^61^, the backbone restraint maintained the E-protein pentamer tertiary structure but the side-chain flexibility in the inner pore was not influenced. A K^+^ as the probe ion, the calculation system was used in the same manner as for umbrella sampling. The ion mean-square displacement (MSD) was calculated along the pore’s Z-axis. A total of 25 simulation systems were obtained, each widows interval distance of the K^+^ being set as 0.24 nm. The umbrella restraint was used to maintain the K^+^ ion’s position on the x-y plane of the pore. The Einstein equation MSD=2D(z)t was used to calculate the diffusion coefficient by the protocol g_msd. The umbrella restraint can be disregarded for these analyses because of the restraint force was negligible compared to thermally induced RMS fluctuations.

#### CE simulations

Computational electrophysiology (CE) is a good tool for simulating the ion conduction ability of a channel under transmembrane voltage. We established a sandwich structure including three parts (membrane-protein-water) as described by Kutzner et al. ^62^ as shown in Fig 4A, each water layer contained a different number of ions. Due to the imbalance of the ion distribution of the water layer, the ions gradient will produce transmembrane voltage. Specific transmembrane voltage can be applied to the simulation system by adjusting the number of ions between the water layers. In this study, we built the SARS-CoV-2 E protein sandwich system. The initial single layer system was equilibrium for 50 ns to make the structures stable. Then the single-layer system was duplicated along the z direction. The resulting system had a size of around 11×11×22 nm^3^. The transmembrane voltage set with different ion number of each parts.

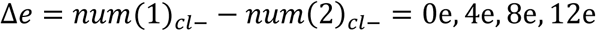

The transmembrane potential was calculated using the implementation of the Poisson equation ∇^2^*U* = −*ρ*/*ε* in the GROMACS g_potential tool. After that, six 20 ns simulations for each transmembrane voltage ware executed, the total simulation time of CE is 6×20×4=480 ns.

The VMD software ^63^ were used to visualize MD trajectories. All of the simulations were performed using the Gromacs 5.1 package ^64^ on Tianhe I Supercomputer in the National Supercomputer Tianjin Center of China. The total simulation times were ∼8 µs.

## Acknowledgments

This study was supported by the National Natural Science Foundation of China (NSFC-31900894 and NSFC-81974464) and The Science & Technology Development Fund of Tianjin Education Commission for Higher Education (2018KJ067). We thank National Supercomputer center – Tianjin provide computing power support.

