## supplemental figures for "Computational Study of Ions and Water Permeation and Transportation Mechanisms of the SARS-CoV-2 Pentameric E Protein Channel"

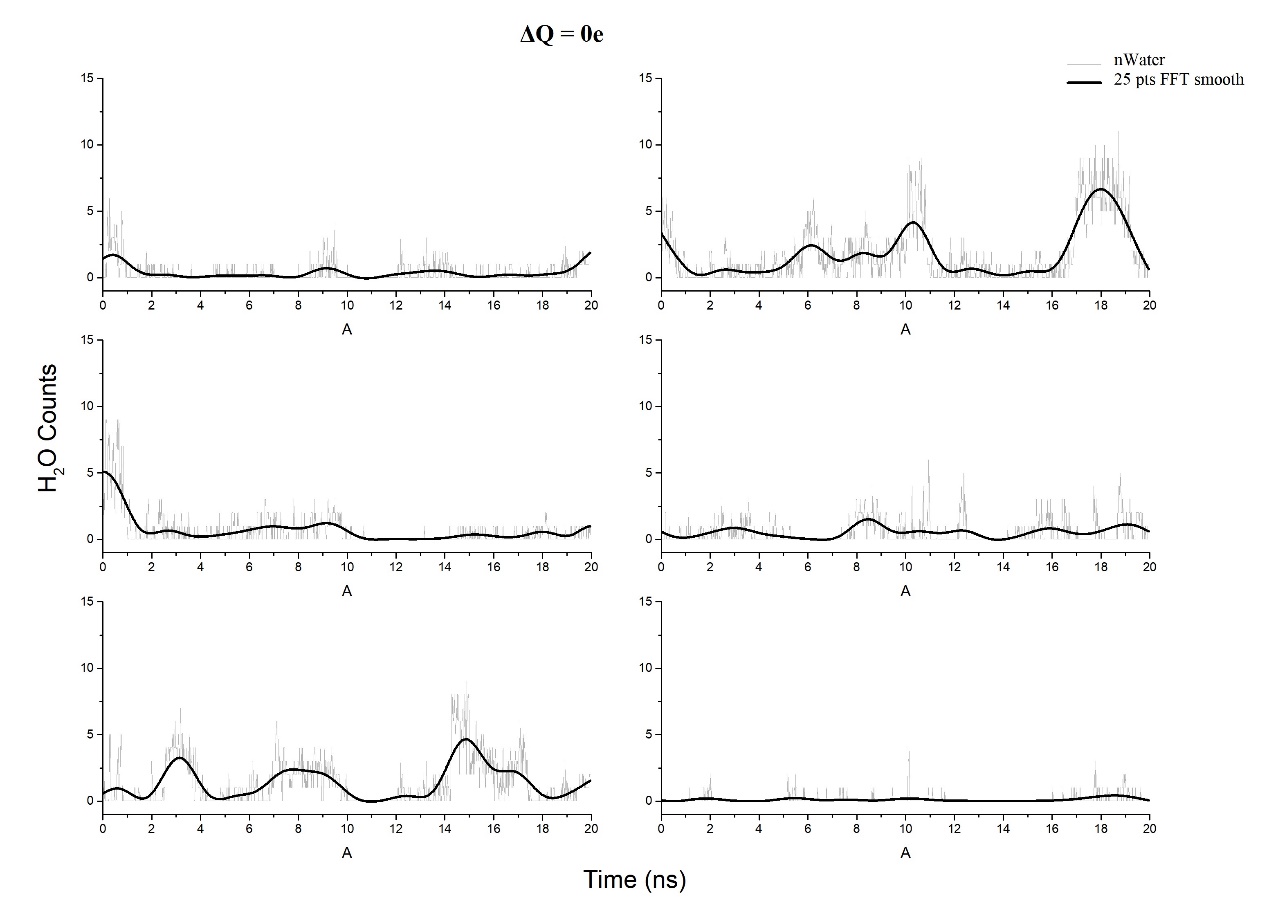

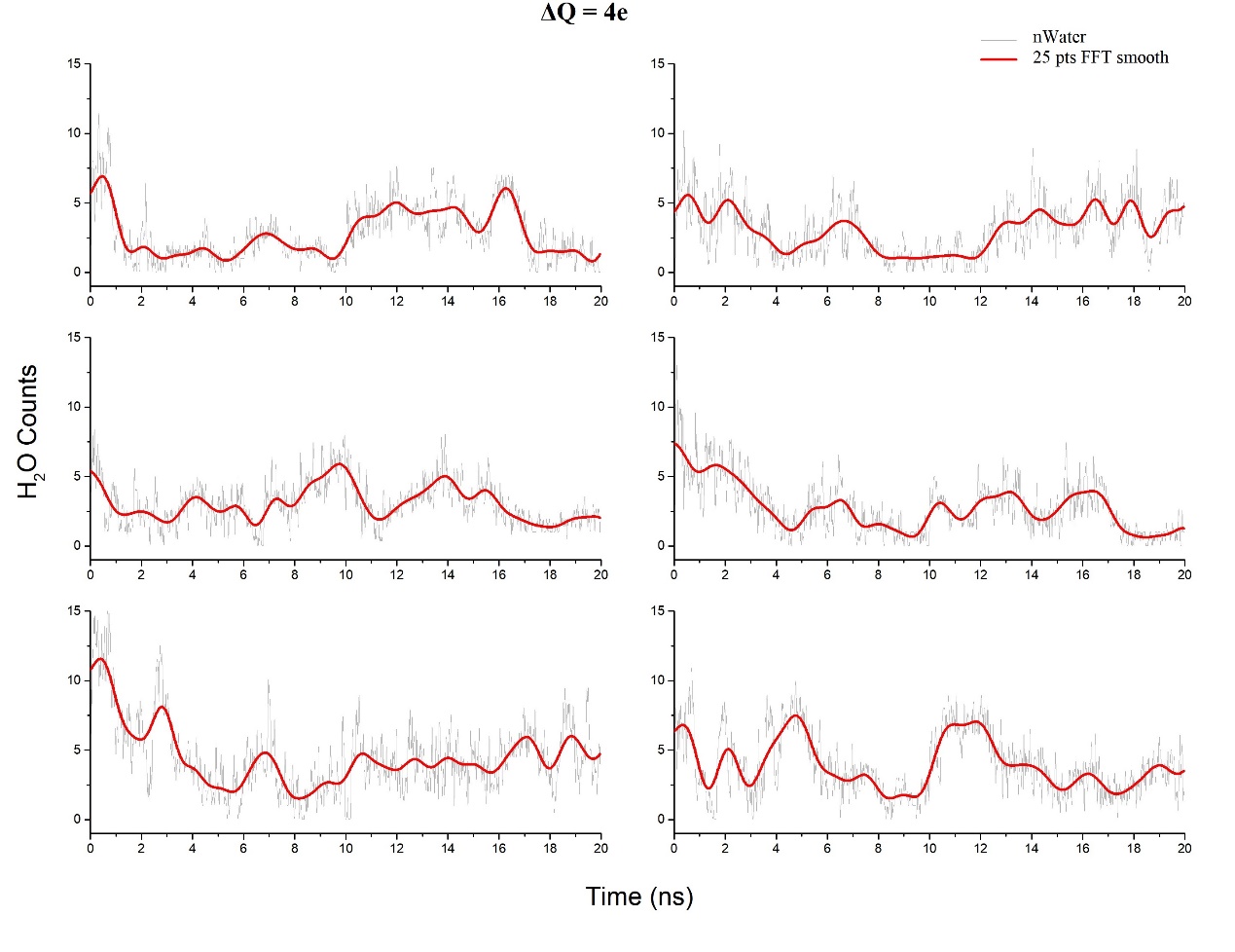

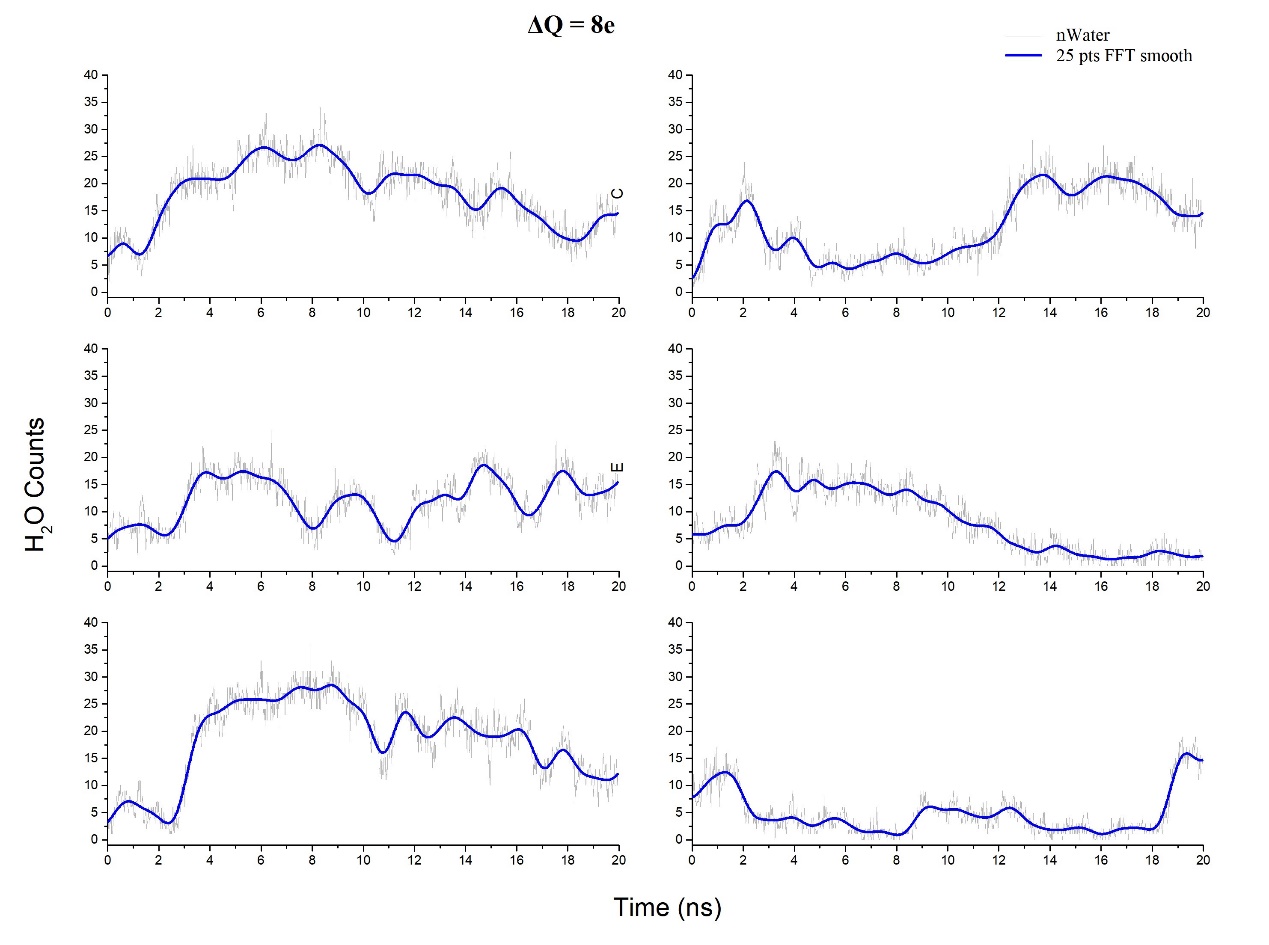

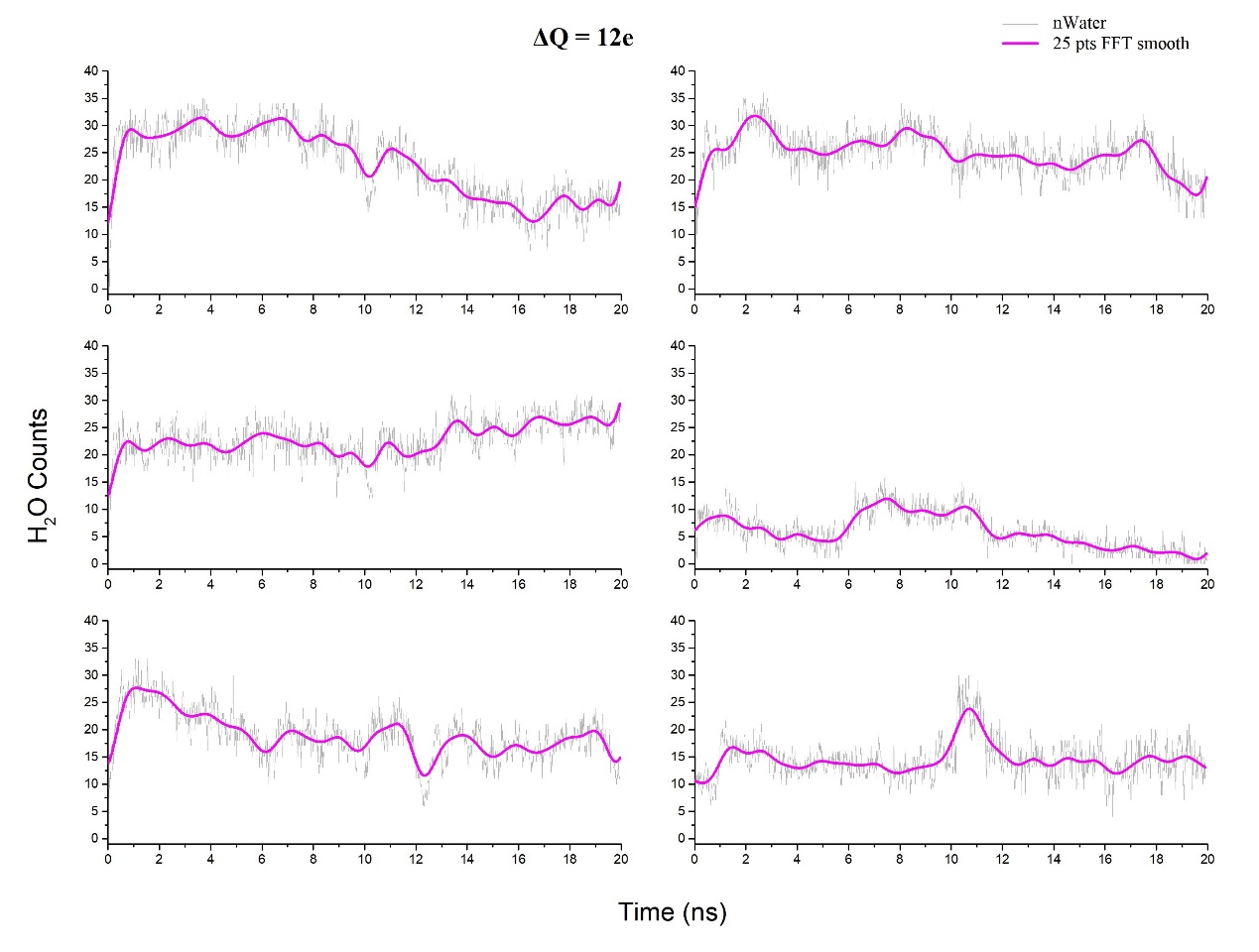

Supplementary Figure. statistical data of the changes in the molecule counts of ΔQ= 0e, 4e, 8e, 12e respectively. The scatter represents the absolute counts of water molecules, and the color curve represents after smoothing.
